# Acute microtubule changes linked to DMD pathology are insufficient to impair contractile function or enhance contraction-induced injury in healthy muscle

**DOI:** 10.1101/2024.06.19.599775

**Authors:** Camilo Vanegas, Jeanine Ursitti, Jacob G. Kallenbach, Kaylie Pinto, Anicca Harriot, Andrew K. Coleman, Guoli Shi, Christopher W. Ward

**Affiliations:** Department of Orthopedics, University of Maryland School of Medicine, Baltimore, MD, USA; Department of Biochemistry and Molecular Biology, University of Maryland School of Medicine, Baltimore, MD, USA; Center for Biomedical Engineering and Technology, University of Maryland School of Medicine, Baltimore, MD, USA

**Keywords:** Duchenne muscular dystrophy, eccentric contraction, Epothilone D, microtubules, skeletal muscle, mechanotransduction

## Abstract

Duchenne muscular dystrophy (DMD) is marked by the genetic deficiency of the dystrophin protein in striated muscle whose consequence is a cascade of cellular changes that predispose the susceptibility to contraction injury central to DMD pathology. Recent evidence identified the proliferation of microtubules enriched in post-translationally modified tubulin as a consequence of dystrophins absence that increases the passive mechanics of the muscle fiber and the excess mechanotransduction elicited reactive oxygen species and calcium signals that promote contraction injury. Motivated by evidence that acutely normalizing the disease microtubule alterations reduced contraction injury in murine DMD muscle (*mdx*), here we sought the direct impact of these microtubule alterations independent of dystrophins absence and the multitude of other changes consequent to dystrophic disease. To this end we used acute pharmacologic (epithiolone-D, EpoD; 4 hours) or genetic (vashohibin-2 and small vasohibin binding protein overexpression via AAV9; 2 weeks) strategies to effectively model the proliferation of detyrosination enriched microtubules in the *mdx* muscle. Quantifying *in vivo* nerve evoked plantarflexor function we find no alteration in peak torque nor contraction kinetics in WT mice modeling these DMD relevant MT alterations. Quantifying the susceptibility to eccentric contraction injury we show EpoD treatment proffered a small but significant protection from contraction injury while VASH/SVBP had no discernable impact. We conclude that the disease dependent MT alterations act in concert with additional cellular changes to predispose contraction injury in DMD.

## Introduction

Duchenne muscular dystrophy (DMD) is a progressive and eventually lethal muscle-wasting disease^1^. Central to DMD pathology is the genetic absence of dystrophin ^2^, a muscle protein serving varied structural and signaling roles^1,3,4^ as an essential linkage between the muscle fiber internal actin and microtubule cytoskeleton and the membrane spanning dystrophin-glycoprotein complex ^1,5^ . Consequent to dystrophin deficiency in the DMD muscle fiber is a progressive cascade of cellular changes that ultimately lead to the decreased muscle specific force production and increased susceptibility to contraction induced injury seen early in disease pathology^1,6^.

Microtubules are dynamic polymers of α- and β-tubulin protein dimers that serve a host of structural, transport and signaling roles in the cell^7^. Central to these microtubule functions are tubulin post-translational modifications (PTM) that regulate MT interaction with protein binding partners including motor proteins, cytoskeletal elements (i.e., actin, intermediate filaments) and dystrophin^5,8,9^. In striated muscle, microtubules enriched in tubulin modified by detyrosination (deTyr-tub) or acetylation (acetyl-tub) positively regulate the mechanics (i.e., stiffness) of the cytoskeleton and the activation of NADPH Oxidase 2 (Nox2) dependent reactive oxygen species (ROS) and calcium (Ca^2+^) signals by mechanotransduction^1–3^. Our works in murine-DMD (*mdx*) show that the proliferation of microtubules enriched in acetyl- and/or deTyr-tub arise as a consequence of the absence of dystrophin and underscore the altered myofibrillar structure^10^, increased muscle fiber mechanics (i.e., stiffness) and the excess mechanotransduction elicited Nox2-ROS and Ca^2+^ signals that underscore contraction injury ^11,12^. Evidence that the acute reduction in the density of microtubules ^12^, or their level of tubulin-PTMs ^11^, is sufficient to prevent contraction injury in murine DMD (*mdx*) established MT alterations as negative disease modifiers and potential therapeutic targets. Subsequent transcriptional ^12^ and proteomic ^13^ evidence of similar tubulin alterations in clinical DMD muscle suggests microtubule pathology is conserved in the human condition. Motivated by the evidence of disease altered microtubules contributing to disease pathology, we sought the direct impact of these microtubule alterations independent from the multitude of other cellular perturbations that arise consequent to dystrophic disease. Given that dystrophic pathology is progressive leading to significant muscle fiber structural alterations and increased fibrosis and fatty infiltrate in the muscle in older *mdx* mice (6-12 months)^14–16^, we benchmarked our disease dependent MT alterations and functional outcomes to those in younger *mdx* mice.

In the present study we challenged 16-week-old wild-type mice with pharmacologic or genetic strategies to model the DMD relevant microtubule alterations in *mdx* mice at this age. Functional measures were then made closely after the pharmacologic (4 hours) or genetic intervention (2 weeks) to realize the DMD relevant microtubule changes while minimizing any longer-term consequences to these microtubule changes. Quantifying nerve evoked plantarflexor function *in vivo* we found that the acute modeling of DMD relevant MT alterations in the WT was insufficient to phenocopy the deficits in isometric torque yet these changes impacted contractile kinetics. Assaying contraction injury, we confirm increased susceptibility to eccentric contraction force loss in the *mdx* yet find a small but significant protection from eccentric contraction induced force loss with EpoD treatment while VASH2/SVBP overexpression has no significant effect. We conclude that consequent to the absence of dystrophin, disease dependent MT alterations act in concert with the myriad of other cellular changes to predispose the force deficits and enhanced contraction injury in DMD muscle.

## Materials and Methods

### Murine models and treatments

All animals were housed and treated in accordance with the standards set by the University of Maryland Baltimore School of Medicine Institutional Animal Care and Use Committee (IACUC). Dystrophic *mdx* (C57BL.10ScSn-Dmdmdx/J) and control (C57BL.10/J) male mice were procured from Jackson Laboratories (Bar Harbor, ME, USA). All mice were housed socially in groups of 3-5 per cage on a 12/12 hr light/dark cycle with food and water *ad libitum*.

Epothilone D (EpoD) is a non-taxane MT targeted clinical chemotherapeutic^17^ that promotes MT polymerization and the accumulation of deTyr and acetyl-tub^18^. Mice were dosed intraperitoneally (IP) with EpoD (10mg/kg) dissolved in dimethylsulfoxide (DMSO; 10mM) or Vehicle (DMSO equal volume to EpoD). Four hours after injection mice were anesthetized for *in vivo* muscle testing and subsequent tissue harvest.

The detyrosination of α-tubulin is by the enzyme complex of vasohibin (VASH) 1 or 2, complexed with small vasohibin binding protein (SVBP). Recent evidence identified VASH2 transcriptionally elevated in young *mdx* muscle vs their WT counterparts. We therefore constructed an AAV9 virus with cDNA encoding VASH2 and SVBP under the muscle-specific promoter MHCK7. The virus was constructed, packaged into AAV9 and ultrapurified by VectorBuilder (vectorbuilder.com). The control and VASH1/SVBP AAV9 used in this study were: pAAV[Exp]-MHCK7>mCherry:WPRE (vector ID: VB211116-1252kch) and pAAV[Exp]-MHCK7>mVash2[NM_144879.2](ns):P2A :mSvbp[NM_001038998.2](ns):P2A:mCherry:WPRE (vector ID: VB211115-1252tbm), respectively. The virus was diluted in sterile saline to 2.5 × 10^11^ GC in 20 μl which was injected in two 10 μl aliquots into the middle of the medial and lateral heads of the gastrocnemius muscle. Control virus was injected in the right leg and VASH2/SVBP virus was injected into the left leg in each of 5 mice. Experiments and tissue collection was performed at 2 weeks post-injection.

### Western blot analyses

Homogenized cell lysates were processed via SDS-PAGE (4-20% BioRad Mini-PROTEAN® TGX™ precast gels), transferred to a membrane (Millipore Immobilon-FL PVDF), stained for total protein (Revert 700 Total Protein Stain, P/N 926-11011, LI-COR Biotechnology) for 5 min at room temperature, then washed in Wash solution (P/N 926-11012, LI-COR Biotechnology) and finally in ultrapure water. Membranes were imaged on a LI-COR Odyssey CLx imager. Immediately after imaging, the membrane was incubated with Revert destaining solution (P/N 926-11013, LI-COR Biotechnology) for 5 minutes and the solution was discarded. The membrane was briefly rinsed in ultrapure water and then blocked with SuperBlock PBS ( 37515; Thermo Scientific) for 1 hr at room temperature. The membrane was probed with primary antibodies overnight for α-Tubulin (2144S, Cell Signaling Technology), β-Tubulin (T4026, Sigma-Aldrich ), detyrosinated Tubulin (31-1335-00, clone RM444, RevMAb Biosciences ) or acetylated Tubulin (T7451, clone 6-11B-1; Sigma-Aldrich ) at 4° C. Blots were washed with 1x TBS + 0.1% Tween 20 (TBST). Blots were then incubated with the corresponding secondary antibody (Li-Cor IRDye® 1:5000) for 1 hour at room temp, washed in TBST and imaged on LI-COR Odyssey CLx imager.

### Immunofluorescence profiling of microtubule structure

Longitudinal cryosections (10-12μm) of snap-frozen extensor digitorum longus (EDL) muscle were air dried to coverslips, fixed with 4% paraformaledehyde in PBS (5 min), washed (3x) in phosphate buffered saline (PBS), then blocked for 15 minutes at room temperature in Superblock blocking buffer in PBS (Thermo Scientific) with 1% Triton™ X-100 (Sigma-Aldrich). Sections were then incubated in primary antibody (β-tubulin; T4026, Sigma-Aldrich) in Superblock PBS at 4° for 72 hours, washed (3x in PBS) then incubated overnight in secondary antibody (goat anti-mouse Alexa Fluor 488; A28175, Invitrogen) at 4°C. Coverslips were then mounted on glass slides using ProLong Gold + DAPI mountant (Invitrogen). Microtubule structure was imaged on a Nikon C2+ confocal fluorescence system coupled to Nikon Ti inverted microscope (20x air objective). Regions of interest (ROI) were manually defined within each myofiber boundary and thresholded to create a binary layer of microtubule structure. The microtubule area was then normalized to the ROI area to calculate microtubule density.

### Muscle performance *in vivo*

Muscle performance was measured *in vivo* with a 305C muscle lever system (Aurora Scientific Inc., Aurora, CAN) as described previously. Briefly, animals were anesthetized in a chamber with 3% isoflurane vapor (SomnoSuite, Kent Scientific) then placed supine on a thermostatically controlled heating pad atop the Aurora 305C with anesthesia maintained via nose cone at ∼2%. The murine hindlimb was secured with a pin at the lateral femoral condyle and the foot was firmly secured to the footplate with cloth tape. Plantarflexor (i..e, gastrocnemius, soleus) contractions were elicited by percutaneous electrical stimulation through the tibial nerve. Optimal isometric twitch force was determined by increasing the current with a minimum of 30 seconds between each twitch to avoid muscle fatigue. Serial electrical stimulations were performed at increasing electrical frequencies of 1, 10, 20, 40, 50, 60, 80, 100 and 150 Hz (0.2 ms pulse width, 500 ms train duration). Following assessment of isometric force, susceptibility to eccentric injury was assayed with 19 eccentric contractions as previously described^11,12,19^. Eccentric contractions were achieved by rotating the footplate 40° backward at a velocity of 800°/s after the first 100 ms of the isometric contraction. The decrease in the peak isometric tetanic force 1 min following the eccentric protocol was taken as the eccentric induced force deficit.

### Statistics

Two-group comparison was performed using t test or Mann–Whitney U test for parametric and nonparametric datasets, respectively. The data are presented as mean ± sem. The only exception is stiffness–indentation velocity relationship curves (Fig. 1, E and G), for which the data comparison was performed using two-way ANOVA, and data are presented as mean ± 95% confidence interval.

**Figure 1.**
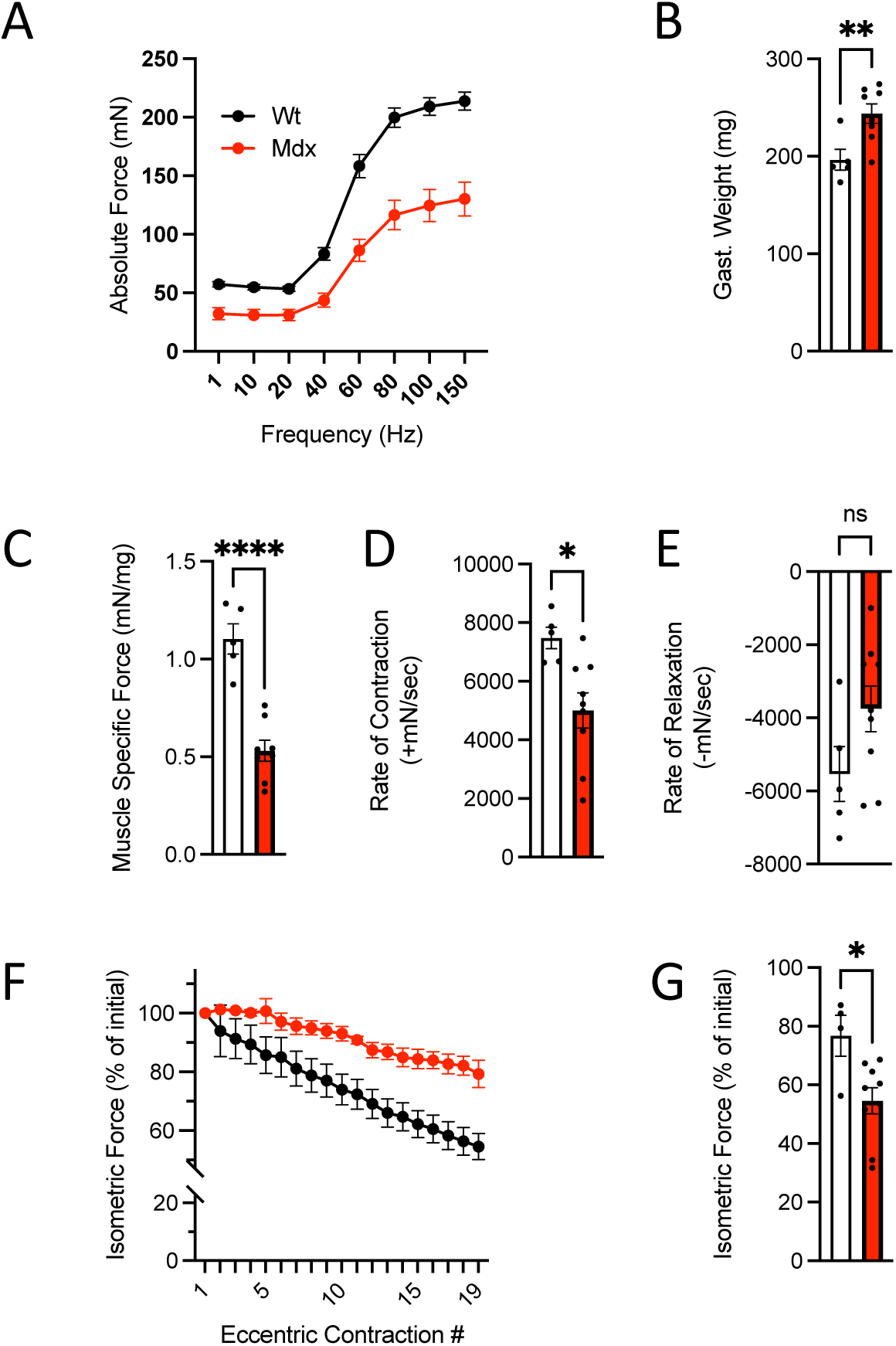
*In vivo* neuromuscular function of C57BL10.*md*x mice (n=9) and C57BL10 controls (n=5). **A.** Force vs stimulation frequency relationship. **B.** Weight of the surgically excised gastrocnemius muscle **C.** Peak isometric force (150Hz) normalized to gastroc mass to yield muscle specific force. The rate of contraction (**D**) and relaxation (**E**) at 150Hz. **F.** Isometric force decline during 19 successive eccentric contractions. **G.** Isometric force 2 min post eccentric contractions.

## Results

Our previous investigations in older *mdx* mice (6-9 months) show that the proliferation of microtubules enriched in acetyl- and/or deTyr-tub arise as a consequence of the absence of dystrophin and underscore the altered myofibrillar structure^10^, increased muscle fiber mechanics (i.e., stiffness) and the excess mechanotransduction-elicited Nox2-ROS and Ca^2+^ signals that underscore contraction injury ^11,12^. Here we sought to determine the direct impact of these disease dependent MT alterations independent of dystrophic disease. To this end, we examined young adult *mdx* mice (13-16 weeks) where pathology is evident, yet the level of secondary pathology within the myofiber (e.g. myofibrillar disorganization) and muscle tissue (e.g., fibrosis) is less advanced and thus less impactful on function^10,14^.

Initial experiments quantified *in vivo* plantarflexor function (i.e., gastrocnemius and soleus) of these young *mdx* and WT mice by examining the isometric force vs stimulation frequency relationship. Here we identified deficits in maximal isometric torque (**Fig 1A**) consistent with previous findings at older ages^9,10^. We also identified increased gastrocnemius mass (**Fig 1B**) in these *mdx* mice, a finding aligned with the pseudohypertrophy early in clinical and murine DMD pathology previously reported^20–22^. Normalizing the isometric torque to gastrocnemius mass (i.e., muscle specific force) realized a further decline in function (**Fig 1C**) which aligns with a decrease in muscle quality contributing to these deficits. Examining the kinetics of muscle contraction, we found the rate-of-contraction significantly impaired in the *mdx*, with no change in the rate of relaxation (**Fig 1D**). These results have been formally attributed to the proliferation of MT’s enriched in deTyr and acetyl-tub increasing the passive mechanics (i.e., stiffness) of the striated muscle fibers^11,23,24^. Finally, we examined the susceptibility of *mdx* muscle to force loss following eccentric contractions; a hallmark of their dystrophic phenotype^25^. Here we find a significant reduction in isometric force in the young *mdx (∼45%) vs.* their WT controls (22%; Fig. **1F-G**). Given reports of older adult *mdx* (6-9 months) losing upwards of ∼75% of their force in this assay^11,12,26–28^, the ∼45% reduction in the young *mdx* was consistent with milder disease pathology at this age.

Our work in mature adult *mdx* mice identified increased tubulin expression and proliferation of MT’s enriched in deTyr-and acetyl-Tub^11,12,24^. Western blot profiling the gastrocnemius muscles from these younger mice also identified elevated tubulin expression (α-Tub) and increased levels of deTyr- and acetyl-Tub in the *mdx* vs WT controls (**Fig 2**). Having established the degree of muscle dysfunction and magnitude of tubulin alterations in the young *mdx*, we sought to model these disease-relevant microtubule alterations in wild-type mice to evaluate their impact on function independent of dystrophic disease.

**Figure 2.**
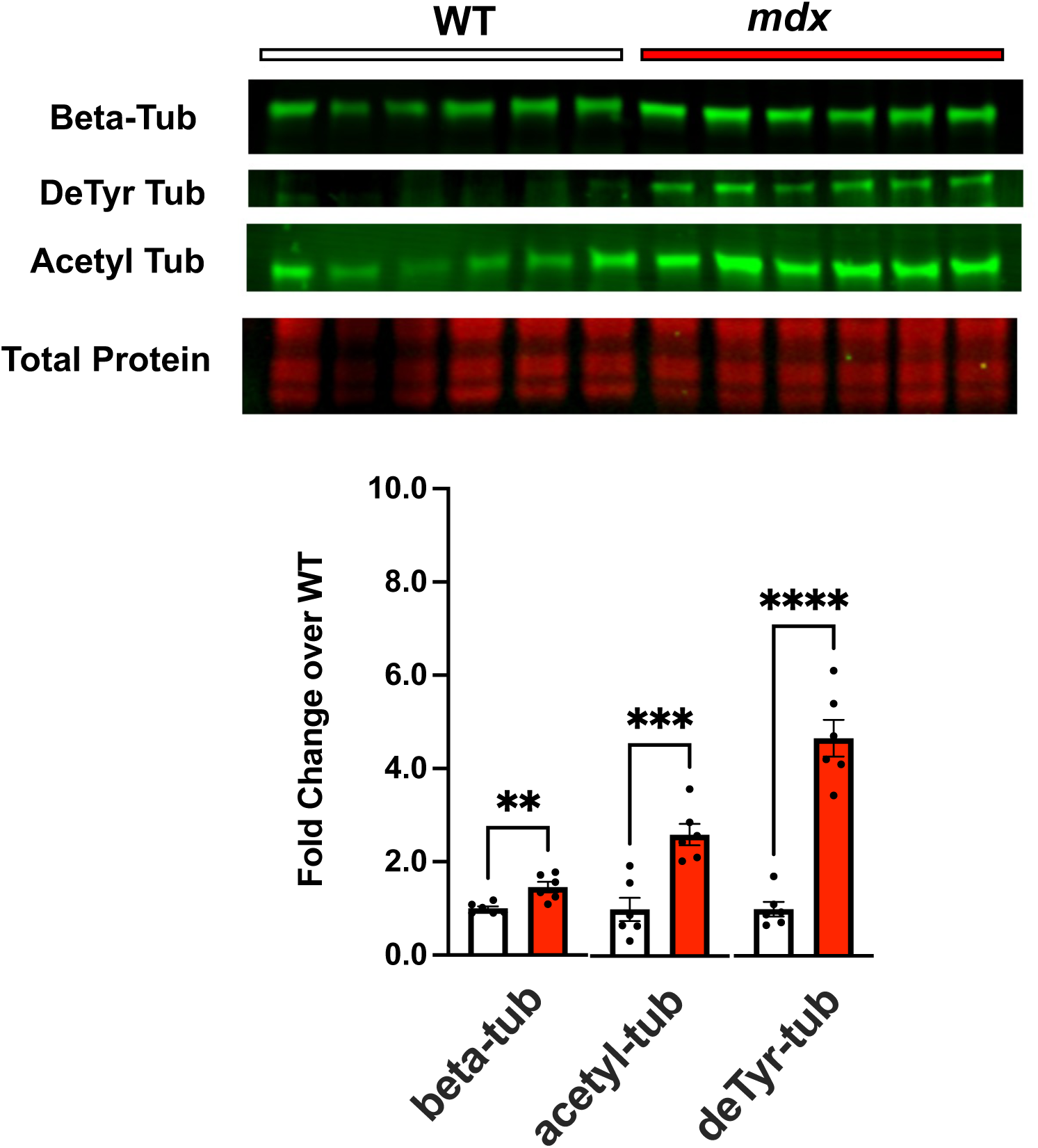
Western blot analysis of gastrocnemius muscle from C57BL10.*md*x mice (n=6) and C57BL10 controls (n=6) probing for levels of tubulin expression (beta tubulin) and tubulins modification by detyrosination and acetylation.

### Acute EpoD treatment models disease relevant microtubule changes in wild-type mice

Young adult C57BL/6J mice were treated with Epothilone D (EpoD) , a tubulin targeted small molecule chemotherapeutic that promotes microtubule polymerization and proliferation and increases the level of deTyr and acetyl-tubulin^20,21^. In brief, mice were dosed with EpoD (10mg/kg; IP) or Vehicle (DMSO; IP) and returned to their home cages. Four hours post-dosing mice were functionally tested, as described above, followed by tissue collection for tubulin biochemistry and histology.

Western blot of the gastrocnemius revealed that EpoD treatment had no impact on tubulin expression, a result consistent with the acute 4-hour timeframe being too brief for significant tubulin expression (**Fig 3**). In contrast we show that this 4-hour exposure effectively increased the level of deTyr-tub (6-fold) and acetyl-tub (1.5-fold) (**Fig 3**) which are levels above or equal to those found in the young adult *mdx* (**Fig 2**). Given that tubulin PTM’s occur on tubulin in the microtubule polymer, we take the evidence of increased deTyr- and acetyl-tub as indirect evidence of microtubule proliferation. We confirmed this by examining microtubule structure from extensor digitorum longus (EDL) muscles isolated from EpoD and DMSO treated animals showing that EpoD treatment significantly increased microtubule proliferation compared to DMSO treated controls (**Fig 4)**.

**Figure 3.**
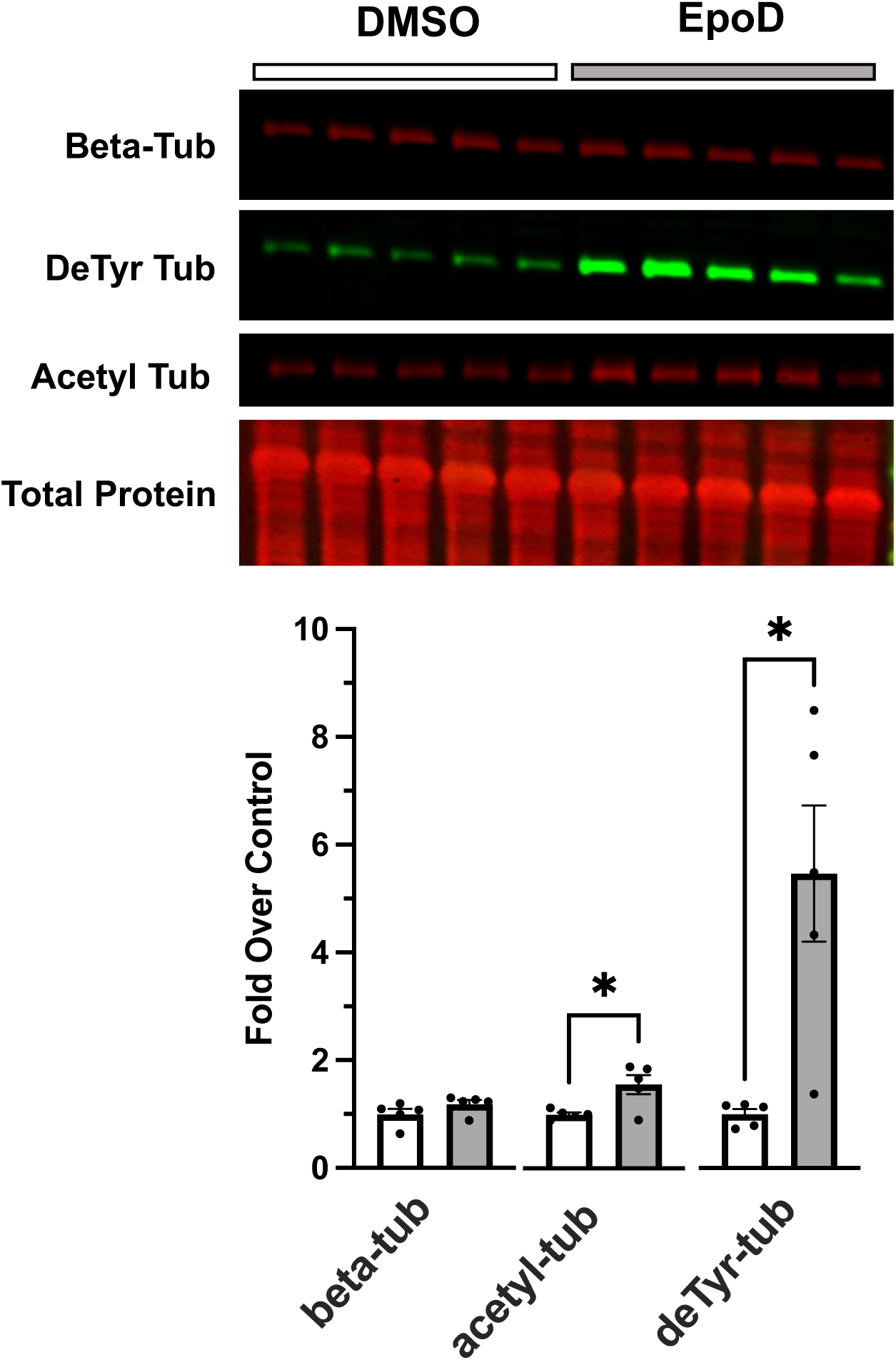
Western blot analysis of gastrocnemius muscle from C57BL10 mice 4 hours post-treatment with either DMSO (control; n=5 ) or EpoD (n=5).

**Figure 4.**
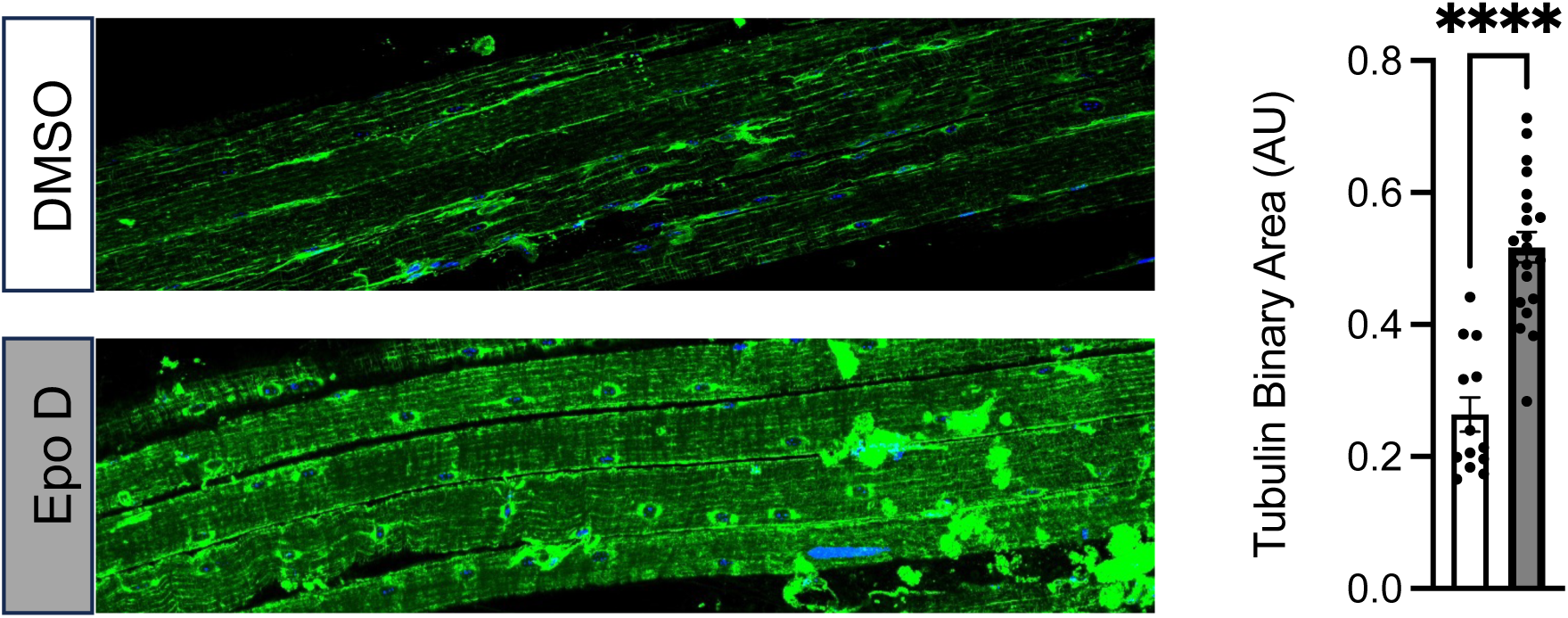
Confocal immunofluorescence images of paraformaldehyde fixed EDL muscle bundles from DMSO or EpoD treated mice labled for beta tubulin. Quantification of beta tubulin pixel area in muscle fibers from the DMSO (n=13) or EpoD (n=19) muscles.

### Acute EpoD treatment has no deleterious impact on neuromuscular function, susceptibility to contraction injury, or passive muscle mechanics

Examining the WT mice treated with either EpoD or vehicle we found no impact on the isometric torque vs stimulation frequency relationship (**Fig 5A**). We also found no impact on the body-weight (not shown) nor the gastrocnemius muscle weight (**Fig 5B**) such that muscle specific force (**Fig 5C**) remained unchanged between treatment groups. Examining the kinetics of the muscle contraction we found no impact on the rate of muscle torque generation (**Fig. 5D**), however EpoD treatment resulted in a slower rate of relaxation (**Fig 5E**). We then challenged mice with eccentric contractions to determine the susceptibility to contraction induced force-loss. Here we showed that EpoD treatment proffered significant protection from eccentric force loss (**Fig 5F**) rather than exacerbating force-loss as we showed in the young *mdx* (Fig 1).

**Figure 5.**
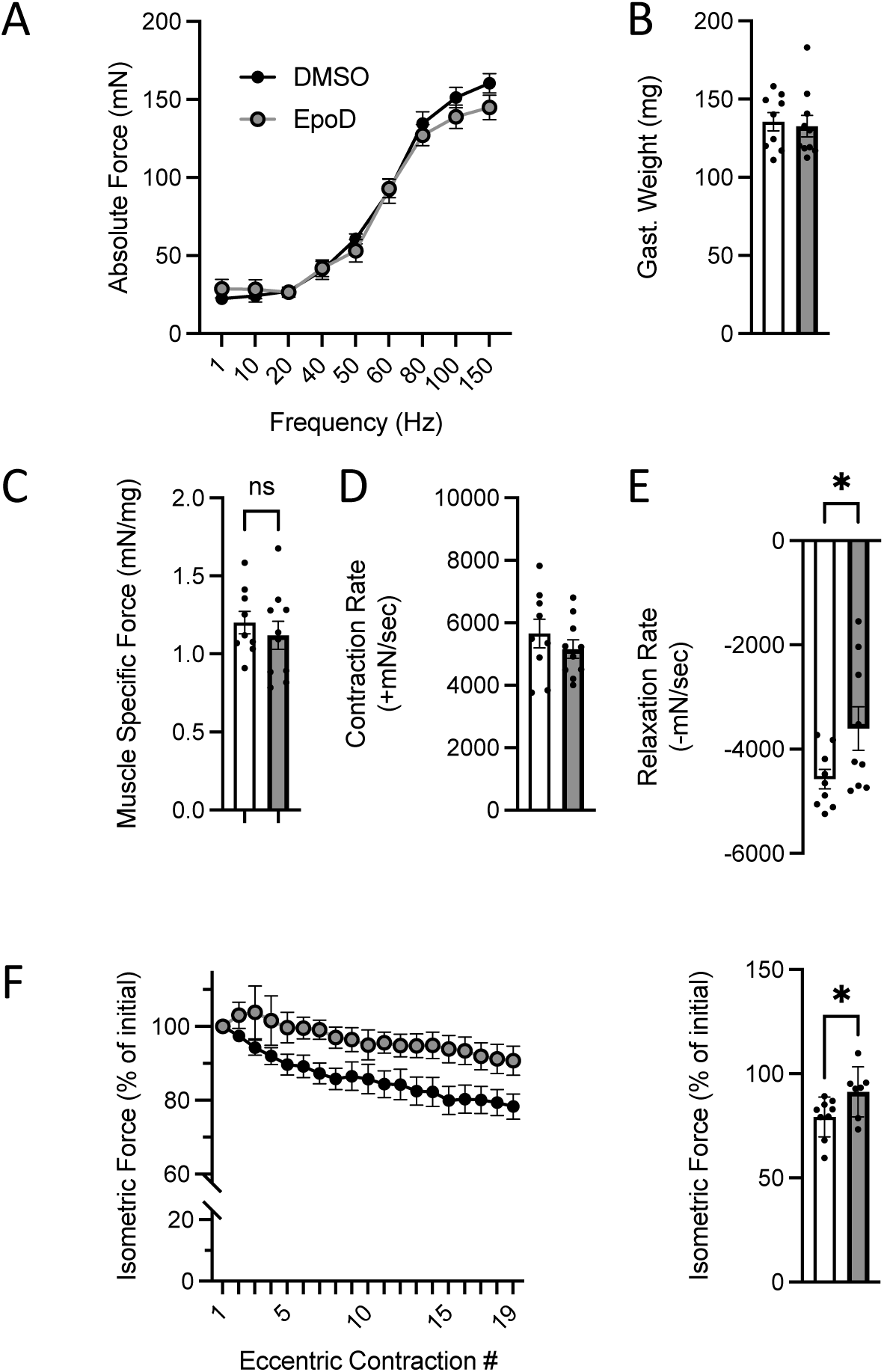
*In vivo* neuromuscular function of C57BL10 mice 4 hours post treatment with either EpoD (n=9) or DMSO (n=9). **A.** Force vs stimulation frequency relationship. **B.** Weight of the surgically excised gastrocnemius muscle **C.** Peak isometric force (150Hz) normalized to gastroc mass to yield muscle specific force. The rate of contraction (**D**) and relaxation (**E**) at 150Hz. **F.** Isometric force decline during 19 successive eccentric contractions. **G.** Isometric force 2 min post eccentric contractions.

### Short-term overexpression of VASH2 and SVBP has no deleterious impact on neuromuscular function or susceptibility to contraction injury

The detyrosination of α-tubulin is by the enzyme complex of vasohibin (VASH) 1 or 2, complexed with small vasohibin binding protein (SVBP)^29,30^. We recently reported that VASH2 transcriptionally elevated in young *mdx* muscle vs their WT counterparts with VASH 1 showing no change^10^. Here we used an AAV9 virus encoding VASH2 and SVBP under control of a skeletal muscle promoter (see methods) delivered by intra-muscular injection to the gastrocnemius muscle of WT mice. AAV9 expression of mCherry under a skeletal muscle promoter in the contralateral gastrocnemius served as the control.

Two weeks post intramuscular injection of AAV9-VASH2/SVBP we find no significant change in the expression of tubulin nor its level of modification by acetylation (i.e., acetyl-tub; **Fig 6**). However, consistent with this enzyme complex being responsible for detyrosination, we find a significant 2.5-fold increase in the level of deTyr-tub. Quantifying the impact of these changes on the isometric torque vs stimulation frequency relationship we find no deleterious impact of the overexpression of VASH2 on the magnitude (**Fig 7A**) nor the kinetics of contraction (**Fig 7D-E**). Muscle mass was not impacted (Fig. 7B) and therefore Muscle Specific Force was unchanged as well (Fig. 7C). Furthermore, we identified no deleterious impact of short-term VASH2/SVBP overexpression on the susceptibility to eccentric contraction induced force-loss (**Fig 7F**).

**Figure 6.**
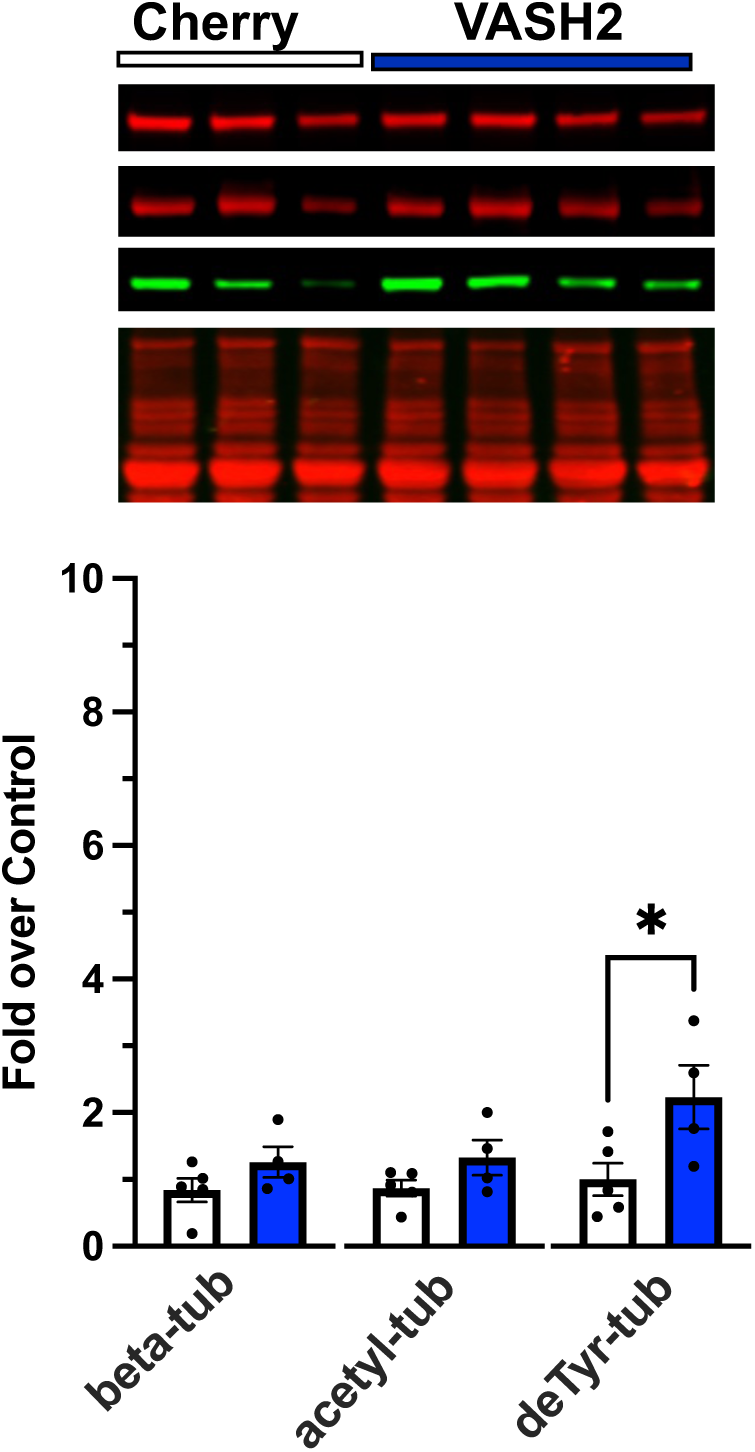
Western blot analysis of gastrocnemius muscle from C57BL10 mice 2 weeks post AAV9 overexpression of either VASH2/SVBP (n=4) or mCherry (control, n=5).

**Figure 7.**
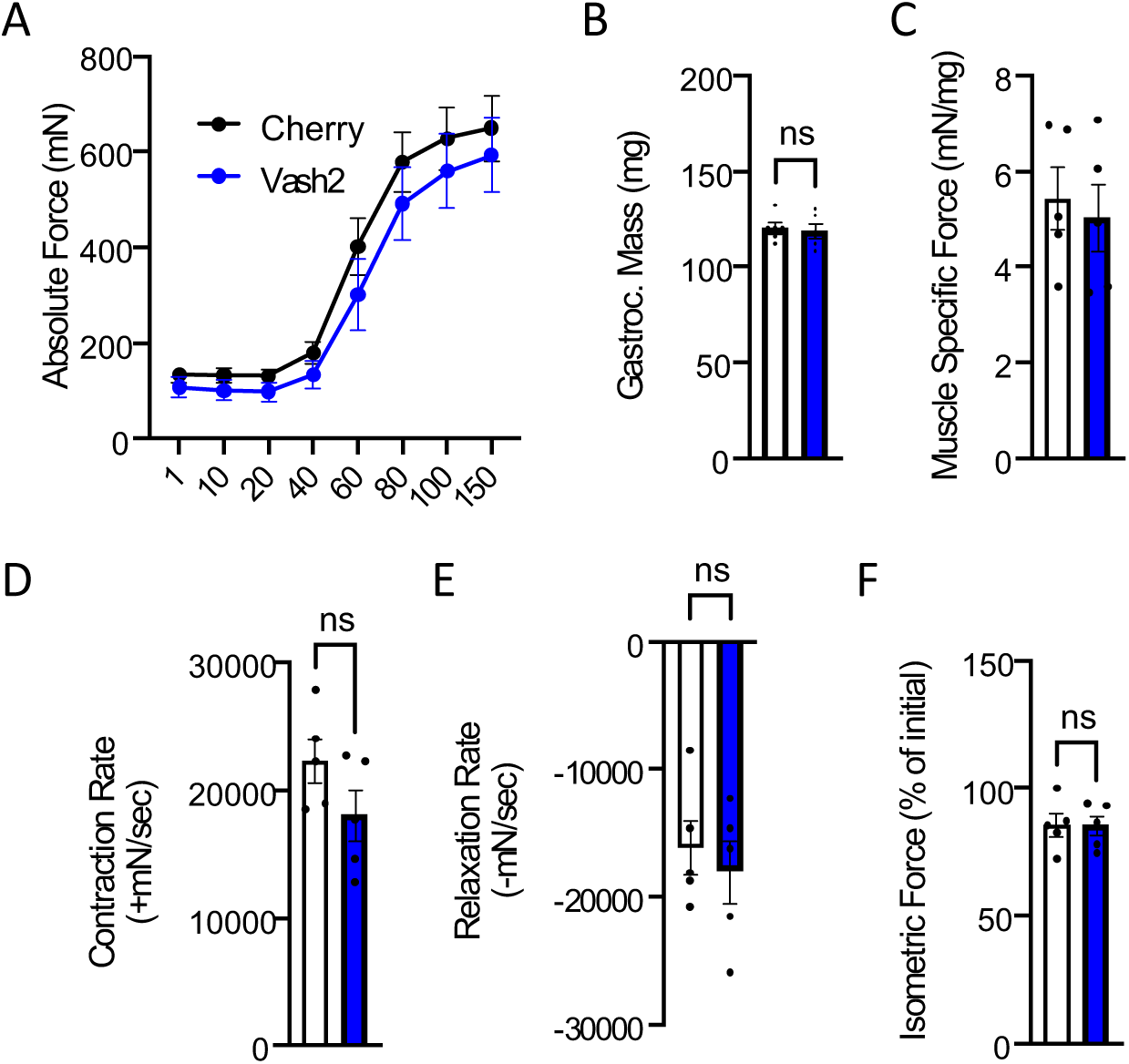
*In vivo* neuromuscular function of C57BL10 mice 2 weeks post AAV9 overexpression of either VASH2/SVBP (n=5) or mCherry (control, n=5) **A.** Force vs stimulation frequency relationship. **B.** Weight of the surgically excised gastrocnemius muscle **C.** Peak isometric force (150Hz) normalized to gastroc mass to yield muscle specific force. The rate of contraction (**D**) and relaxation (**E**) at 150Hz. **F.** Isometric force 2 min post eccentric contractions.

## Discussion

The genetic absence of dystrophin elicits a progressive cascade of signaling and structural changes in skeletal muscle that underscore the deficits in muscle force and susceptibility to contraction injury central to dystrophic pathology^1^. Consistent with dystrophins role as a cytolinker to microtubules at the sub-sarcolemmal membrane, dystrophins absence predisposes a disorganized sub-sarcolemmal microtubule network that becomes highly proliferated as disease progresses^5,32^. Our initial discoveries found these disease proliferated microtubules enriched in tubulin modified by acetylation ( acetyl-tub) and/or detyrosination (deTyr-tub) which increase the passive mechanics (i.e., stiffness) of the muscle fiber and the magnitude of Nox2-ROS and Ca^2+^ signaling by mechanotransduction^11,12,24^. We recently expanded the consequence of these microtubule alterations by linking the proliferation of deTyr-tub enriched microtubules to the altered myofibrillar structure in *mdx* muscle fibers^10^.

Consistent with these microtubule associated signaling and structural alterations as negative disease modifiers we showed that the acute pharmacologic reduction in microtubule abundance^12^, or the level of deTyr-tub^11^, in the *mdx* mouse was sufficient to decrease the dystrophic muscles susceptibility to contraction injury. While evidence of transcript and proteomic enrichment of deTyr-tub in muscle of DMD boys ^12^ ^13^ supports the clinical relevance of these microtubule changes, evidence they occur in parallel to other diverse cellular changes (i.e., fibrosis, altered myofibrillar structure, mitochondrial dysfunction) motivated our interest to dissect the direct impact of the microtubule alterations independent of disease.

Here we show that the acute modeling of DMD relevant MT changes in WT muscle is insufficient to recapitulate the impaired contractile function or enhanced susceptibility to contraction-induced injury seen in the *mdx* mouse. Given this result we conclude that the disease relevant MT alterations act in concert with other disease dependent alterations to yield functional deficits.

One consequence of dystrophic pathology is increased expression of Nox2 complex proteins, which together with the microtubule changes, drive the deleterious mechanotransduction elicited Nox2-ROS and Ca^2+^ signals that contribute to contraction induced force loss^11,12,33,34^. Evidence that targeting either Nox2 or microtubules effectively reduces deleterious Nox2-ROS and Ca^2+^ signaling and contraction force loss^11,12,33,34^ suggests both microtubule changes and a threshold level of oxidative stress may be necessary elements to realize muscle dysfunction. Consistent with this concept is evidence that EpoD has minimal side effects with short term dosing^18^ but enhances pathology in mice with reduced oxidative buffering capacity (i.e., SOD1 null)^35^. Further insight comes from our result that EpoD treatment enhanced, rather than diminished, the ability to sustain isometric force during a brief bout of successive eccentric contractions. This result aligns with evidence that physiologic levels of mechano-elicited Nox2-ROS regulate Ca^2+^ influx^36^ and metabolic pathways^37,38^ necessary to sustain repetitive muscle activation. Motivated by these findings our future work will explore the potential synergy of disease relevant oxidative stress and microtubule alterations in contributing to contraction induced force loss.

## Acknowledgements

This work was supported by the National Institutes of Health grants R01-AR071618 and R01-AR071614 (to C.W.W) and 2T32-AR007592-26 (to J.G.K).

## Conflicts of Interest

Christopher W. Ward is the Chief Scientific Officer of Myologica, LLC. All other authors declare no conflicts of interest.

## Author Contributions

J. Ursitti, C. Vanegas, J.G. Kallenbach, and C.W. Ward designed the experiments. Western blot analyses were conducted by C. Vanegas, A.K. Coleman, and G. Shi. Immunofluorescent imaging and quantification of myofibers were conducted by A. Harriot and K. Pinto. Muscle physiology was conducted by C. Vanegas and J.G. Kallenbach. Results were analyzed and interpreted by C. Vanegas, J.G. Kallenbach, and C. W. Ward. The manuscript was written by J. Ursitti, C. Ward, Vanegas, J.G. Kallenbach, and C.W. Ward. All authors reviewed, edited, and finalized the manuscript.

## Notes

### Competing Interest Statement

The authors have declared no competing interest.

## References

1. Khairallah, R. J. et al. Microtubules underlie dysfunction in duchenne muscular dystrophy. Sci. Signal. 5, ra56–ra56 (2012).

2. Kerr, J. P. et al. Detyrosinated microtubules modulate mechanotransduction in heart and skeletal muscle. Nat. Commun. 6, 1–14 (2015).

3. Coleman, A. K., Joca, H. C., Shi, G., Lederer, W. J. & Ward, C. W. Tubulin acetylation increases cytoskeletal stiffness to regulate mechanotransduction in striated muscle. J. Gen. Physiol. 153, e202012743 (2021).

4. Prins, K. W. et al. Dystrophin is a microtubule-associated protein. J. Cell Biol. 186, 363–369 (2009).

5. Liu, W. & Ralston, E. A new directionality tool for assessing microtubule pattern alterations. Cytoskeleton 71, 230–240 (2014).

6. Oddoux, S. et al. Misplaced Golgi Elements Produce Randomly Oriented Microtubules and Aberrant Cortical Arrays of Microtubules in Dystrophic Skeletal Muscle Fibers. Front. Cell Dev. Biol. 7, 1–18 (2019).

7. Belanto, J. J. et al. Independent variability of microtubule perturbations associated with dystrophinopathy. Hum Mol Genet 25, 4951–4961 (2016).

8. Prosser, B. L., Khairallah, R. J., Ziman, A. P., Ward, C. W. & Lederer, W. J. X-ROS signaling in the heart and skeletal muscle: stretch-dependent local ROS regulates [Ca^2+^]i. J. Mol. Cell. Cardiol. 58, 172–81 (2013).

9. De Silva S, Fan Z, Kang B, Shanahan CM, Zhang Q. Nesprin-1: novel regulator of striated muscle nuclear positioning and mechanotransduction. Biochem Soc Trans. 2023 Jun 28;51(3):1331–1345. doi: 10.1042/BST202215411. Allen, D. G., Whitehead, N. P. & Froehner, S. C. Absence of Dystrophin Disrupts Skeletal Muscle Signaling: Roles of Ca2+, Reactive Oxygen Species, and Nitric Oxide in the Development of Muscular Dystrophy. Physiol Rev 96, 253–305 (2016).

2. Hoffman, E. P., Brown, R. H. & Kunkel, L. M. Dystrophin: the protein product of the Duchenne muscular dystrophy locus. Cell 51, 919–928 (1987).

3. Constantin, B. Dystrophin complex functions as a scaffold for signalling proteins. Biochim Biophys Acta 1838, 635–42 (2014).

4. Li, D., Yue, Y., Lai, Y., Hakim, C. H. & Duan, D. Nitrosative stress elicited by nNOSmicro delocalization inhibits muscle force in dystrophin-null mice. J Pathol 223, 88–98 (2011).

5. Prins, K. W. et al. Dystrophin is a microtubule-associated protein. J. Cell Biol. 186, 363–369 (2009).

6. Allen, D. G., Zhang, B. & Whitehead, N. P. Stretch-Induced Membrane Damage in Muscle: Comparison of Wild-Type and mdx Mice. in Muscle Biophysics: From Molecules to Cells (ed. Rassier, D. E.) 297–313 (Springer, New York, NY, 2010). doi:10.1007/978-1-4419-6366-6_17.

7. Akhmanova, A. & Lukas C Kapitein. Mechanisms of microtubule organization in differentiated animal cells. Nat. Rev. Mol. Cell Biol. (2022) doi:10.1038/s41580-022-00473-y.

8. Janke, C. & Magiera, M. M. The tubulin code and its role in controlling microtubule properties and functions. Nat. Rev. Mol. Cell Biol. (2020) doi:10.1038/s41580-020-0214-3.

9. Roll-Mecak, A. The Tubulin Code in Microtubule Dynamics and Information Encoding. Dev. Cell 54, 7–20 (2020).

10. Harriot, A. D. et al. Detyrosinated microtubule arrays drive myofibrillar malformations in mdx muscle fibers. Front. Cell Dev. Biol. 11, 1209542 (2023).

11. Kerr, J. P. et al. Detyrosinated microtubules modulate mechanotransduction in heart and skeletal muscle. Nat. Commun. 6, 8526 (2015).

12. Khairallah, R. J. et al. Microtubules underlie dysfunction in duchenne muscular dystrophy. Sci Signal 5, ra56 (2012).

13. Capitanio, D. et al. Comparative proteomic analyses of Duchenne muscular dystrophy and Becker muscular dystrophy muscles: changes contributing to preserve muscle function in Becker muscular dystrophy patients. J. Cachexia Sarcopenia Muscle 11, 547–563 (2020).

14. Massopust, R. T. et al. Lifetime analysis of mdx skeletal muscle reveals a progressive pathology that leads to myofiber loss. Sci. Rep. 10, 17248 (2020).

15. Friedrich, O. et al. Microarchitecture is severely compromised but motor protein function is preserved in dystrophic mdx skeletal muscle. Biophys J 98, 606–16 (2010).

16. Head, S. I. Branched fibres in old dystrophic mdx muscle are associated with mechanical weakening of the sarcolemma, abnormal Ca2+ transients and a breakdown of Ca2+ homeostasis during fatigue. Exp Physiol 95, 641–56 (2010).

17. Vahdat, L. T. Clinical Studies With Epothilones for the Treatment of Metastatic Breast Cancer. Semin. Oncol. 35, S22–S30 (2008).

18. Altaha, R., Fojo, T., Reed, E. & Abraham, J. Epothilones: A Novel Class of Non-taxane Microtubule-stabilizing Agents. Curr. Pharm. Des. 8, 1707–1712 (2002).

19. Boyer, J. G. et al. Depletion of skeletal muscle satellite cells attenuates pathology in muscular dystrophy. Nat. Commun. 13, 2940 (2022).

20. Vohra, R. S. et al. Magnetic Resonance Assessment of Hypertrophic and Pseudo-Hypertrophic Changes in Lower Leg Muscles of Boys with Duchenne Muscular Dystrophy and Their Relationship to Functional Measurements. PLoS One 10, e0128915 (2015).

21. Froehner, S. C., Reed, S. M., Anderson, K. N., Huang, P. L. & Percival, J. M. Loss of nNOS inhibits compensatory muscle hypertrophy and exacerbates inflammation and eccentric contraction-induced damage in mdx mice. Hum Mol Genet 24, 492–505 (2015).

22. Faber, R. M., Hall, J. K., Chamberlain, J. S. & Banks, G. B. Myofiber branching rather than myofiber hyperplasia contributes to muscle hypertrophy in mdx mice. Skelet Muscle 4, 10 (2014).

23. Robison, P. et al. Detyrosinated microtubules buckle and bear load in contracting cardiomyocytes. Science 352, aaf0659 (2016).

24. Coleman, A. K., Joca, H. C., Shi, G., Lederer, W. J. & Ward, C. W. Tubulin acetylation increases cytoskeletal stiffness to regulate mechanotransduction in striated muscle. J. Gen. Physiol. 153, e202012743 (2021).

25. Lovering, R. M. & De Deyne, P. G. Contractile function, sarcolemma integrity, and the loss of dystrophin after skeletal muscle eccentric contraction-induced injury. Am J Physiol Cell Physiol 286, C230–8 (2004).

26. Hakim, C. H., Grange, R. W. & Duan, D. The passive mechanical properties of the extensor digitorum longus muscle are compromised in 2- to 20-mo-old mdx mice. J Appl Physiol 1985 110, 1656–63 (2011).

27. Baltgalvis, K. A. et al. Transgenic overexpression of gamma-cytoplasmic actin protects against eccentric contraction-induced force loss in mdx mice. Skelet Muscle 1, 32 (2011).

28. Baumann, C. W., Ingalls, C. P. & Lowe, D. A. Mechanisms of weakness in Mdx muscle following in vivo eccentric contractions. J. Muscle Res. Cell Motil. 43, 63–72 (2022).

29. Aillaud, C. et al. Vasohibins/SVBP are tubulin carboxypeptidases (TCPs) that regulate neuron differentiation. Science 358, 1448–1453 (2017).

30. Nieuwenhuis, J. et al. Vasohibins encode tubulin detyrosinating activity. Science 358, 1453–1456 (2017).

31. Olson, M. T. et al. Taurine Is Covalently Incorporated into Alpha-Tubulin. J. Proteome Res. 19, 3184–3190 (2020).

32. Belanto, J. J. et al. Independent variability of microtubule perturbations associated with dystrophinopathy. Hum Mol Genet 25, 4951–4961 (2016).

33. Loehr, J. A. et al. NADPH oxidase mediates microtubule alterations and diaphragm dysfunction in dystrophic mice. Elife 7, (2018).

34. Loehr, J. A. et al. Eliminating Nox2 reactive oxygen species production protects dystrophic skeletal muscle from pathological calcium influx assessed in vivo by manganese-enhanced magnetic resonance imaging. J Physiol 594, 6395–6405 (2016).

35. Clark, J. A. et al. Epothilone D accelerates disease progression in the SOD1 ^G93A^ mouse model of amyotrophic lateral sclerosis. Neuropathol. Appl. Neurobiol. 44, 590–605 (2018).

36. Michaelson, L. P., Iler, C. & Ward, C. W. ROS and RNS signaling in skeletal muscle: critical signals and therapeutic targets. Annu Rev Nurs Res 31, 367–87 (2013).

37. Bedard, K. & Krause, K. H. The NOX family of ROS-generating NADPH oxidases: physiology and pathophysiology. Physiol Rev 87, 245–313 (2007).

38. Brandes, R. P., Weissmann, N. & Schroder, K. Nox family NADPH oxidases in mechano-transduction: mechanisms and consequences. Antioxid Redox Signal 20, 887–98 (2014).

